# 12-Lipoxygenase inhibition delays onset of autoimmune diabetes in human gene replacement mice

**DOI:** 10.1101/2024.07.28.604986

**Authors:** Titli Nargis, Charanya Muralidharan, Jacob R. Enriquez, Jiayi E. Wang, Kerim Kaylan, Advaita Chakraborty, Sarida Pratuangtham, Kayla Figatner, Jennifer B. Nelson, Sarah C. May, Jerry L. Nadler, Matthew B. Boxer, David J. Maloney, Sarah A. Tersey, Raghavendra G. Mirmira

**Affiliations:** Department of Medicine and the Kovler Diabetes Center, The University of Chicago, Chicago, IL 60637, USA; Department of Pharmacology, New York Medical College, Valhalla, NY 10595, USA; Veralox Therapeutics, Frederick, MD 21704, USA

**Keywords:** type 1 diabetes, islet, macrophage, autoimmunity, lipoxygenase, mouse model

## Abstract

Type 1 diabetes (T1D) is characterized by the autoimmune destruction of insulin-producing β cells and involves an interplay between β cells and cells of the innate and adaptive immune systems. We investigated the therapeutic potential of targeting 12-lipoxygenase (12-LOX), an enzyme implicated in inflammatory pathways in β cells and macrophages, using a mouse model in which the endogenous mouse *Alox15* gene is replaced by the human *ALOX12* gene. Our finding demonstrated that VLX-1005, a potent 12-LOX inhibitor, effectively delayed the onset of autoimmune diabetes in human gene replacement non-obese diabetic mice. By spatial proteomics analysis, VLX-1005 treatment resulted in marked reductions in infiltrating T and B cells and macrophages with accompanying increases in immune checkpoint molecule PD-L1, suggesting a shift towards an immune-suppressive microenvironment. RNA sequencing analysis of isolated islets and polarized proinflammatory macrophages revealed significant alteration of cytokine-responsive pathways and a reduction in interferon response after VLX-1005 treatment. Our studies demonstrated that the *ALOX12* human replacement gene mouse provides a platform for the preclinical evaluation of LOX inhibitors and supports VLX-1005 as an inhibitor of human 12-LOX that engages the enzymatic target and alters the inflammatory phenotypes of islets and macrophages to promote the delay of autoimmune diabetes.

## INTRODUCTION

The pathogenesis of type 1 diabetes (T1D) involves a complex interplay between multiple cell types within the pancreatic islet, including innate immune cells (macrophages, dendritic cells), insulin-producing cells (β cells), and adaptive immune cells (T cells, B cells) (1). Although the disease has traditionally been viewed as arising from a primary defect in immune tolerance, an emerging perspective posits that environmental factors (such as viruses or other systemic inflammatory disorders) may aggravate an interaction between macrophages and β cells, facilitating oxidative and endoplasmic reticulum (ER) stress pathways in β cells (2–4). These pathways facilitate the generation of β-cell neoepitopes that then trigger adaptive autoimmunity (5, 6). Disease-modifying therapies—those that alter disease pathogenesis rather than correcting the underlying disease phenotypes—have largely focused on the adaptive immune system and seen some successes in clinical trials. For example, an anti-CD3 monoclonal antibody (teplizumab) that targets activated T cells has been shown to delay the onset of T1D by up to two years in subjects at high risk for the disease (7). Given the increasing appreciation of innate immune cells and β cells in early T1D pathogenesis, the identification of drugs targeting these cell types raises the possibility that combination therapeutic approaches may provide more durable outcomes.

The lipoxygenases (LOXs) encompass a family of enzymes involved in lipid metabolism that facilitates the oxygenation of polyunsaturated fatty acids to form eicosanoids, some of which are pro-inflammatory in nature (8). In the mouse, 12/15-LOX is encoded by the *Alox15* gene and is the primary active LOX present in macrophages and β cells and produces the proinflammatory eicosanoid 12-hydroxyeicosatetraenoic acid (12-HETE) as a principal product from the substrate arachidonic acid (9). Whole-body deletion of *Alox15* on the autoimmune non-obese diabetic (NOD) mouse background results in almost complete protection against diabetes (10). Deletion of *Alox15* in either the innate immune myeloid cells (2) or in β cells (11) recapitulates the autoimmune diabetes protection seen in the whole-body deletion, emphasizing both the early role of these cell types in T1D and the importance of the 12/15-LOX pathway in disease pathogenesis. In these cell-specific deletion models, islets exhibit marked reductions in invading pathogenic T cells (insulitis), a finding reflecting the disease-modifying response. The molecular events tied to disease protection ostensibly emanate from reductions in oxidative and ER stress (and the resultant reduction in neoepitope formation and presentation) as well as from enhanced display of PD-L1 (an immune-suppressive checkpoint ligand) on the surface of myeloid cells and β cells (2, 11).

In humans, the relevant LOX enzyme that produces 12-HETE is 12-LOX, encoded by the *ALOX12* gene. Like the mouse 12/15-LOX, human 12-LOX is present in residual insulin-positive cells in donors with T1D or in autoantibody-positive donors at risk for T1D (12)—a finding consistent with a potential role in promoting β-cell sensitivity to autoimmunity. A major challenge to using mice as a platform to test inhibitors is that human 12-LOX exhibits structurally distinct characteristics from mouse 12/15-LOX, thereby necessitating the development of different inhibitors that cannot be tested for efficacy in mice (13–15). Previously, VLX-1005 (also known as ML355) was described as a potent and selective inhibitor of human 12-LOX while also displaying a favorable half-maximal inhibitory concentration (IC_50_) and pharmacokinetic (PK) properties (16). VLX-1005 was shown to protect human islets in vitro against dysfunction caused by proinflammatory cytokines (17), but the lack of appropriate in vivo model systems has made it challenging to pharmacologically validate VLX-1005 as a therapeutic target in autoimmune diabetes. To address this challenge, we developed new mouse strains in which the mouse *Alox15* gene is replaced by the human *ALOX12* gene while retaining the mouse gene’s upstream control elements. This human gene replacement platform was leveraged to test if and how human 12-LOX pharmacologic inhibition of human 12-LOX with VLX-1005 modifies disease progression in autoimmune T1D.

## RESULTS

### Generation and validation of the hALOX12 gene replacement mouse model

To establish a platform to test potential inhibitors of human 12-LOX in vivo, we generated a mouse model in which the endogenous mouse *Alox15* gene is replaced by the human *ALOX12* gene (**Figure 1A**). This model leaves the mouse upstream regulatory region intact to ensure that the expression of *ALOX12* recapitulates the expression of *Alox15*. These mice (henceforth referred to as *hALOX12* mice) were introgressed onto the *C57BL/6J* mouse background using a speed congenics approach and bred to homozygosity. Microsatellite genotyping showed that the mice were 100% congenic on the *C57BL/6J* (or simply *B6*) background (**Supplemental Table 1** Excel file). To confirm the successful deletion of mouse *Alox15* and replacement with human *ALOX12*, we performed standard genotyping (**Supplemental Figure 1A**). Additionally, we isolated tissues (kidney, spleen, lung, fat, liver, islets, peritoneal macrophages, and bone marrow-derived macrophages (BMDMs)) from wildtype *B6* and *B6.hALOX12* mice and subjected them to gene expression analysis for *Alox15* and *ALOX12.* As anticipated, wildtype tissues expressed mouse *Alox15* and did not express human *ALOX12*; conversely, *B6.hALOX12* mice tissues expressed *ALOX12* but not *Alox15* (**Table 1**).

**Figure 1:**
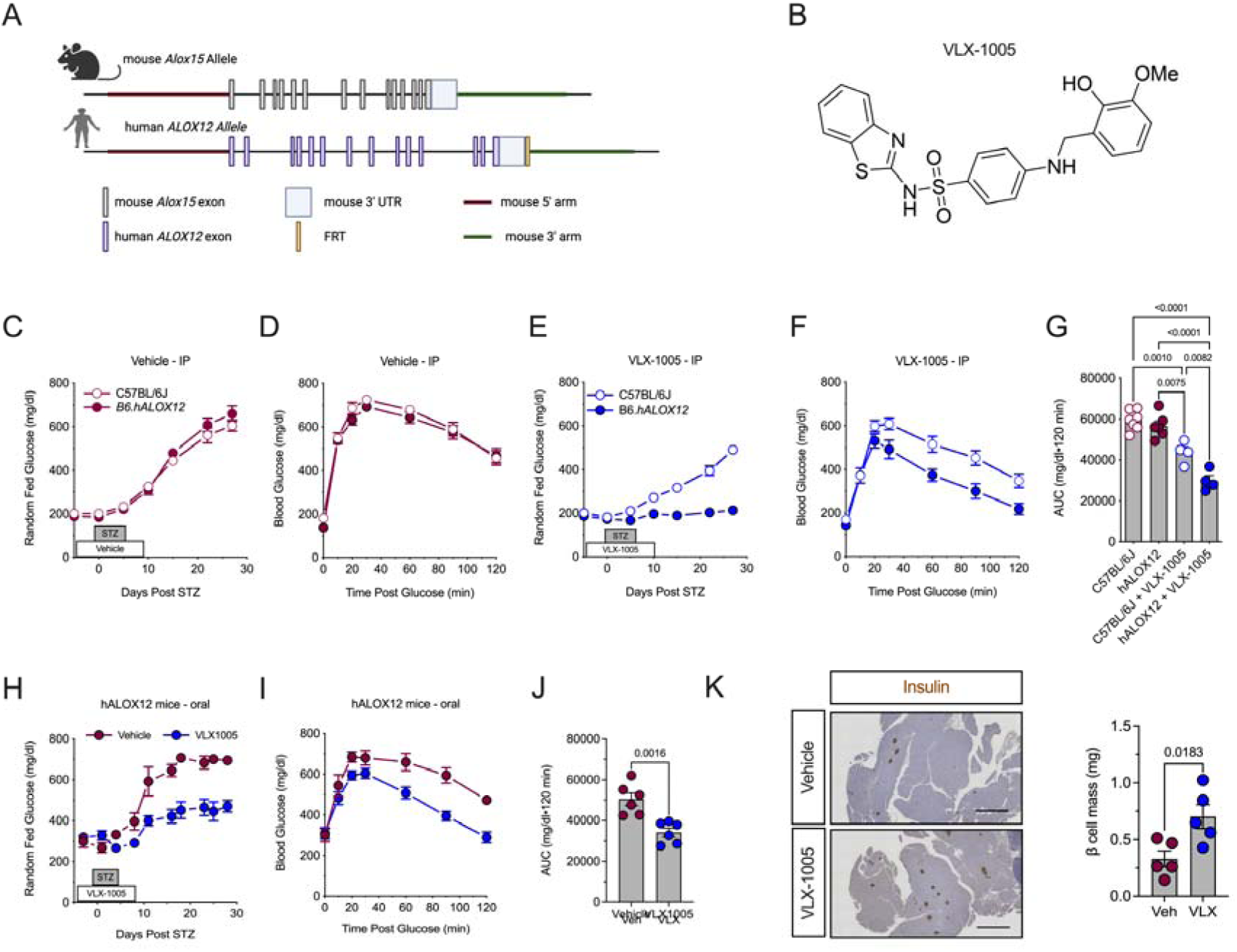
12-LOX inhibition protects against streptozotocin-induced diabetes. *C57BL/6J* and *B6*.*hALOX12* male mice (N=4-7 per group as indicated) were treated with 30 mg/kg intraperitoneal or PO VLX-1005 and multiple low-dose streptozotocin (STZ). (**A**) Schematic of the generation of *hALOX12* mice by replacing mouse *Alox15* with human *ALOX12*. (**B**) Chemical structure of VLX-1005. (**C**) Random-fed blood glucose values in vehicle-treated male *C57BL/6J* and *B6*.*hALOX12* mice after STZ. (**D**) GTT of vehicle-treated male *C57BL/6J* and *B6*.*hALOX12* mice after STZ at day 4 post-STZ-treatment. (**E**) Random-fed blood glucose values in VLX-1005-treated male *C57BL/6J* and *B6*.*hALOX12* mice after STZ. (**F**) GTT of VLX-1005-treated male *C57BL/6J* and *B6*.*hALOX12* mice after STZ at day 4 post-STZ-treatment. (**G**) AUC of *C57BL/6J* and *B6*.*hALOX12* after STZ at day 4 post-STZ-treatment (one-way ANOVA). (**H**) Random-fed blood glucose values in male vehicle-or VLX-1005-treated (PO) *B6*.*hALOX12* mice after STZ. (**I**) GTT of male vehicle- or VLX-1005-treated (PO) *B6*.*hALOX12* mice after STZ at day 4 post-STZ-treatment. (**J**) AUC of *B6*.*hALOX12* after STZ at day 4 post-STZ-treatment. (**K**) Pancreata stained for insulin (left panel) and β cell mass measurement (right panel) from male *B6*.*hALOX12* mice at day 26 post-STZ-treatment. Scale bars = 500 μm. Data are presented as mean ±SEM and statistical significance was determined by a two-tailed T-test or one-way ANOVA.

**Table 1:**
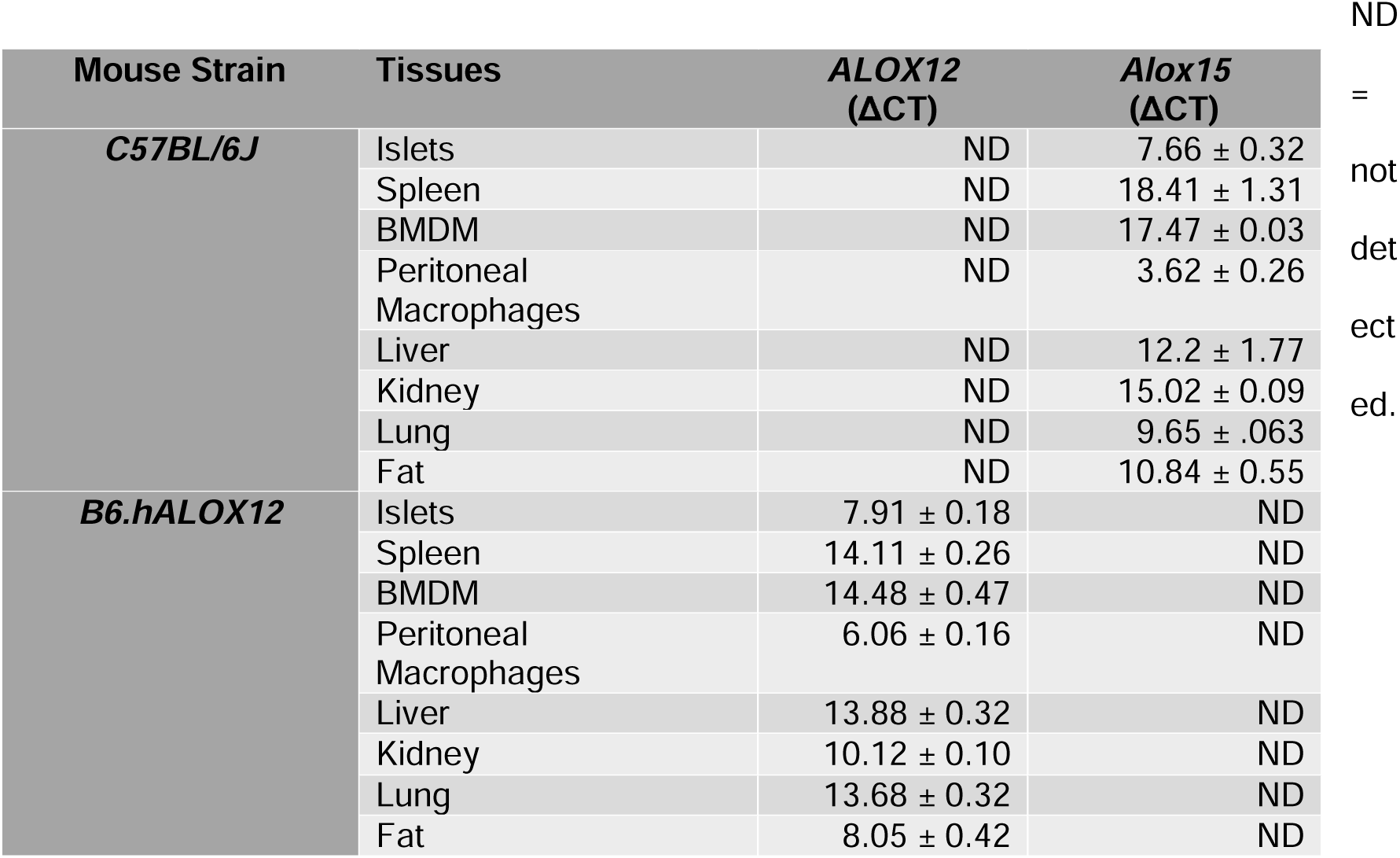
RNA expression levels of human *ALOX12* and mouse *Alox15* normalized to mouse *Actb* from various isolated tissues of *C57BL/6J* and *B6*.*hALOX12* mice.

Because lipoxygenases are known to affect metabolic function, we next performed metabolic characterization to determine if/how the replacement of *Alox15* with *ALOX12* altered metabolic phenotypes. We found no significant differences in body weight, lean mass, fat mass, random-fed blood glucose levels, or glucose tolerance between wildtype *B6* and *B6.hALOX12* mice (**Supplemental Figure 1B-F**). Moreover, islet ultrastructure (relative immunostaining patterns of α cells and β cells) and composition (α cell mass and β cell mass) were indistinguishable between 10 week-old wildtype and *B6.hALOX12* mice (**Supplemental Figure** 1G-I). Taken together, these data suggest that the successful replacement of *Alox15* with human *ALOX12* did not alter gross metabolic or islet phenotypes.

### Effects of VLX-1005 against STZ-induced diabetes are specific to B6.hALOX12 mice

Prior studies demonstrated that whole-body deletion of mouse *Alox15* protects against diabetes induced by the chemical streptozotocin (STZ) (18). To test if the human 12-LOX inhibitor VLX-1005 (14) (**Figure 1B**) phenocopies deletion of the enzyme in our human gene replacement mice, we employed a similar STZ diabetes induction protocol. STZ is a β cell toxin that induces low-grade inflammation, macrophage influx into islets, and eventual diabetes in mice after 5 daily low-dose intraperitoneal injections (55 mg/kg) (19). Eight-week-old male wildtype *B6* and *B6.hALOX12* mice were injected intraperitoneally daily with vehicle or 30 mg/kg VLX-1005 in the peri-STZ treatment period (for the 5 days before, during, and after STZ). STZ-injected *B6* and *B6.hALOX12* mice receiving vehicle became overtly hyperglycemic within 10 days of starting STZ treatment and displayed equivalent glucose intolerance by GTT (**Figure 1C and D**). Upon receiving VLX-1005, however, *B6.hALOX12* mice showed complete protection from STZ-induced diabetes, whereas wildtype *B6* mice became overtly hyperglycemic (**Figure 1E**); GTTs at the end of the study confirmed improved glucose tolerance in *B6.hALOX12* mice compared to wildtype *B6* mice (**Figure 1F and G**). These data indicate a specific effect of the drug in preventing hyperglycemia in *B6.hALOX12* mice and support the effectiveness of the *hALOX12* platform for interrogating VLX-1005 action.

### Pharmacokinetics of oral VLX-1005 and its effects on STZ-induced diabetes

Given that the oral route is the preferred route for systemic drug delivery in humans, we next asked if oral administration of VLX-1005 provides adequate exposure in mice. We performed pharmacokinetic analysis following a single oral administration (as a suspension in 0.5% methylcellulose) of VLX-1005 spray dried dispersion at a dose of 30 mg/kg in C57BL/6J mice, followed by serial analysis of VLX-1005 levels by LC-MS/MS. The pharmacokinetic profile of orally-administered VLX-1005 in mice shows a mean half-life (T_1/2_) of 3.24 ± 0.07 hours and a consistent T_max_ of 0.250 hours across all mice. The C_max_ was 13300 ± 624 ng/ml, with moderate variability in AUC (15029 ± 3177 h*ng/ml). These parameters, particularly the low variability in T_max_ and C_max_, support the feasibility of once-daily dosing for maintaining therapeutic levels over a 24-hour period (**Table 2**). We next tested the effects of oral administration of VLX-1005 on the low-dose STZ model, with VLX-1005 (at 30 mg/kg) given 3 days prior to the start of STZ, during STZ, and for 3 days following STZ treatment. Similar to intraperitoneal delivery, oral administration of VLX-1005 in *B6.hALOX12* mice resulted in lower random-fed blood glucose levels (**Figure 1H**) and significantly improved glucose tolerance (**Figure 1I and J**) compared to vehicle—although this effect was not as robust as with intraperitoneal delivery of the drug. Consistent with improved glucose homeostasis, oral VLX-1005-treated mice exhibited greater β cell mass at the end of the study compared to vehicle-treated mice (**Figure 1K**). Collectively, these data suggest that a single daily oral delivery of VLX-1005 (at 30 mg/kg) achieves plasma levels with therapeutic efficacy.

**Table 2:** Plasma concentration vs time profile for VLX-1005 after 30 mg/kg PO in *C57BL/6J* mice and *NOD.ShiLt/J* mice.

| Pharmacokinetic Parameter | C57BL/6J |  | NOD.ShiLt/J |  |
| --- | --- | --- | --- | --- |
|  | Mean ± SD | CV (%) | Mean ± SD | CV (%) |
| *T<sub>1/2</sub> (h) | 3.24 ± 0.07 | 2.17 | 2.53 ± 0.41 | 16.1 |
| T<sub>max</sub> (h) | 0.250 ± 0.00 | 0.00 | 0.250 ± 0.00 | 0.000 |
| C<sub>max</sub> (ng/ml) | 13300 ± 624 | 4.70 | 14253 ± 5474 | 38.4 |
| AUC<sub>last</sub> (h*ng/ml) | 15029 ± 3177 | 21.1 | 13211 ± 1631 | 12.3 |
| AUC<sub>Inf</sub> (h*ng/ml) | 15083 ± 3206 | 21.3 | 13225 ± 1625 | 12.3 |
| AUC<sub>_%Extrap_obs</sub> (%) | 0.342 ± 0.133 | 38.9 | 0.115 ± 0.099 | 85.6 |
| MRT<sub>Inf_obs</sub> (h) | 2.57 ± 0.38 | 14.8 | 3.81 ± 0.71 | 18.7 |
| AUC<sub>last/D</sub> (h*ng/ml) | 501 ± 106 | 21.1 | 440 ± 54 | 12.3 |

\*T<sub>1/2</sub>, half life; T<sub>max</sub>, time to maximum drug concentration; C<sub>max</sub>, maximum drug concentration; AUC<sub>last</sub>, area under the curve from the time of dosing to the last measurable concentration; AUC<sub>Inf</sub>, area under the curve from the time of dosing extrapolated to infinity; AUC<sub>\_%Extrap\_obs</sub>, area under the curve from the time of dosing extrapolated to last observed concentration; MRT<sub>Inf\_obs</sub>, mean residence time from the time of dosing extrapolated to infinity; AUC<sub>last/D</sub>, dose normalized area under the curve to time of last measurable concentration.

### VLX-1005 treatment reduces ***β*** cell inflammation in NOD.hALOX12 mice

The non-obese diabetic (*NOD*) mouse model is a model of T1D that recapitulates many of the immune and β cell features of the disease (20). We, therefore, asked if pharmacologic inhibition of 12-LOX using orally administered VLX-1005 protects against spontaneous diabetes development in the *NOD* mouse model. To address this question, we introgressed humanized *hALOX12* mice onto the *NOD* background using a speed congenics approach. Genome scanning of microsatellites was performed to confirm that mice were 100% congenic on the *NOD* mouse background (*NOD.hALOX12* mice) (**Supplemental Table 1** Excel file). We next measured human 12-LOX protein levels in the *NOD.hALOX12* mice. Similar to the gene profile we observed in *B6.ALOX12 mice* compared to wildtype *C57BL/6J* mice, wildtype *NOD* tissues robustly expressed mouse 12/15-LOX (the protein encoded by mouse *Alox15*) and little/no human *ALOX12*; conversely, *NOD.hALOX12* mice tissues robustly expressed human 12-LOX and minimal levels of 12/15-LOX (**Supplemental Figure 1J and 1K**). Consistent with their congenic nature, female *NOD.hALOX12* mice exhibited islet pathology similar to *NOD* mice at the (prediabetic) age of 10 weeks with evidence of T and B cell infiltration of islets (**Supplemental Figure 1L**) and indistinguishable insulitis score (**Supplemental Figure 1M)**, suggesting that replacement of *Alox15* with human *ALOX12* did not alter the islet pathology of the disease. Pharmacokinetics of orally administered VLX-1005 (30 mg/kg) on the *NOD* background were similar to those seen in *C57BL/6J* mice (**Table 2**), suggesting that the *NOD* background does not affect drug absorption or clearance.

To assess the effect of VLX-1005 administration on products of 12-LOX activity in *NOD.hALOX12* mice, we administered 30 mg/kg VLX-1005 (or vehicle) orally to female *NOD.hALOX12* mice for 1 week during the pre-diabetic phase (8 weeks of age) and harvested serum. Lipidomics analysis (by LC/MS/MS) was performed for a series of 12-LOX products resulting from different fatty acid substrates (**Figure 2A**). Notably, levels of 12-HETE (from arachidonic acid), 13-HODE (from linoleic acid), and 14-HDHA and 17-HDHA (from docosahexaenoic acid) were all significantly reduced (**Figure 2B**). Levels of 12-HEPE (from eicosapentaenoic acid) were not significantly changed (**Figure 2B**), suggesting minimal involvement of eicosapentaenoic acid metabolism in *NOD* mice. Lipids within the pathway that are processed by enzymes other than 12-LOX were not statistically significantly altered (**Supplemental Table 2**). These data are collectively consistent with the expected 12-LOX engagement by VLX-1005.

**Figure 2:**
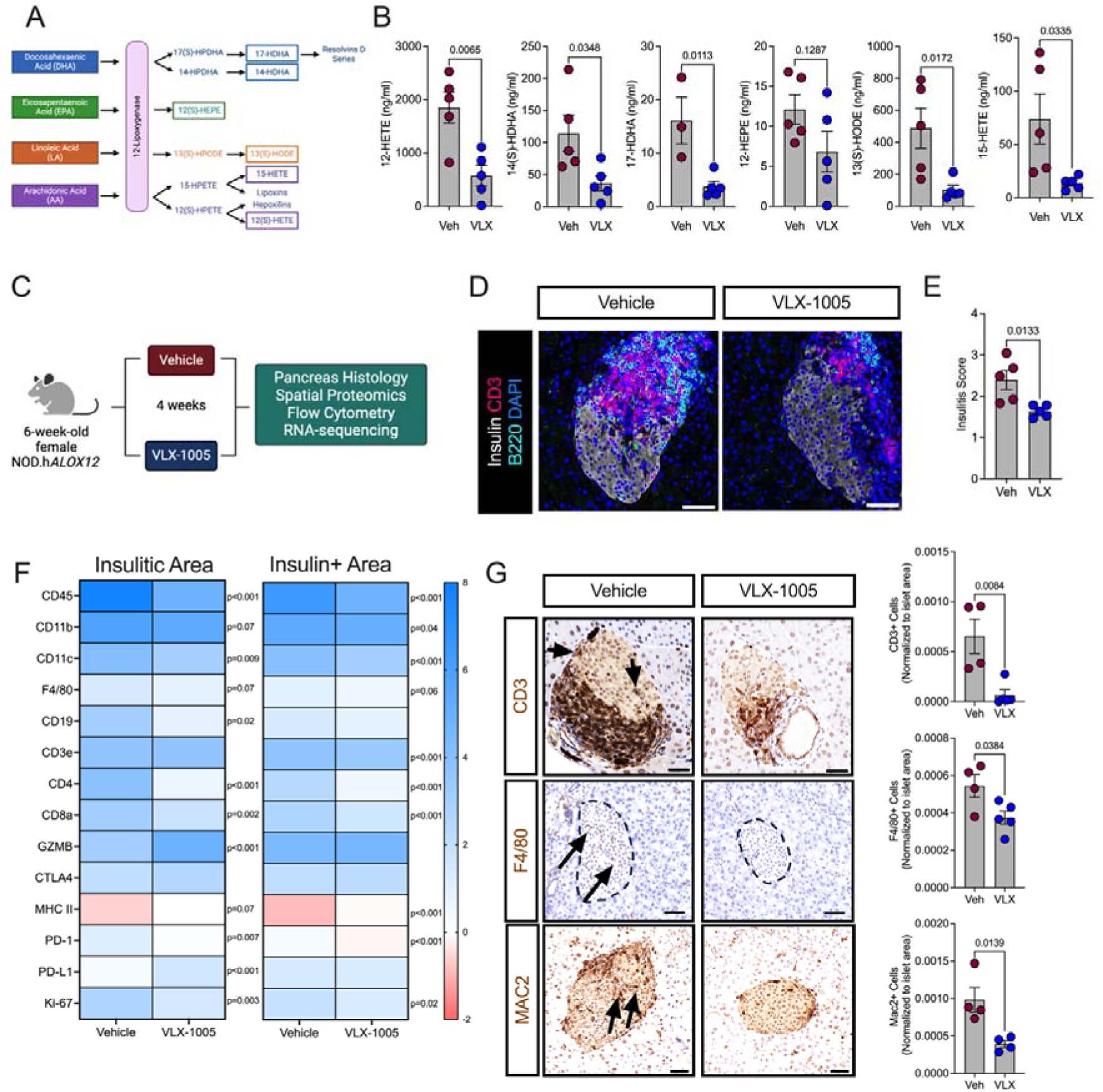
VLX-1005 decreased islet inflammation in *NOD*.*hALOX12* female mice. 6-week-old female pre-diabetic *NOD*.*ALOX12* mice were treated orally with 30 mg/kg VLX-1005 for 4 weeks prior to tissue analysis. (**A**) Schematic representation of 12-lipoxygenase products. (**B**) Serum lipidomics results of 12-lipoxygnease products as indicated (N=4-5). (**C**) Schematic representation of mouse treatment paradigm. (**D**) Pancreata from mice stained for CD3 (magenta), B220 (teal), insulin (white), and nuclei (blue). Scale bars = 50 μm. (**E**) Average insulitis score, each dot represents an individual mouse (N=4-5). (**F**) Heatmap of identified proteins in the insulitic area (*left panel*) and insulin-positive area (*right panel*). (**G**) Pancreata of mice stained and quantified for CD3 (brown, *top panels*; *arrows* indicate positive CD3 staining within the islet), F4/80 (brown, *middle panels*; *arrows* indicate positive F4/80 staining within the islet), or MAC2 (brown, *bottom panels*; *arrows* indicate positive MAC2 staining within the islet) and nuclei (blue). Each dot represents an individual mouse (N=4-5).Scale bars = 50 μm. Data are presented as mean ±SEM and statistical significance was determined by a two-tailed T-test in all cases.

To assess the effect of VLX-1005 administration on immune cell phenotypes in *NOD.hALOX12* mice, we administered 30 mg/kg VLX-1005 (or vehicle) orally to female *NOD.hALOX12* mice for 4 weeks during the pre-diabetic phase (6-10 weeks of age) and harvested pancreas, pancreatic lymph nodes (pLNs), and spleen (**Figure 2C**). Pancreas pathology showed reduced T and B cell infiltration and that the extent of insulitis (by insulitis scoring) was significantly reduced in VLX-1005-treated *NOD.hALOX12* mice compared to vehicle-treated mice (**Figure 2D-E**). To specifically interrogate the nature of immune cells within the insulitic region, we performed spatial tissue-based proteomics (Nanostring®). We used insulin immunostaining and nuclei staining to identify β cells and the surrounding insulitic regions. Pre-validated antibodies in the GeoMx® mouse immune panel were used to probe for immune cell subtypes in the peri-islet insulitic region and within the islet. *NOD.hALOX12* mice exhibited a notable reduction of myeloid population subtypes in both insulitic and islet areas, including macrophages (F4/80+; CD11b+) and dendritic cells (CD11c+) (**Figure 2F** and **Supplemental Figure 2A-B**). This reduction in myeloid cell populations was accompanied by a decrease in T and B cells populations, including CD4+, CD3+, CD8+, and CD19+ cells (**Figure 2F** and **Supplemental Figure 2A-B**). Immunohistochemistry of pancreas sections confirmed the reductions in both T cells (CD3+), macrophages (F4/80+), and activated macrophages (Mac2+) following oral VLX-1005 treatment (**Figure 2G**). A notable observation in spatial proteomics was the increased levels of the immune checkpoint ligand PD-L1 (**Figure 2F**). Enhanced PD-L1/PD-1 interactions shift T cells to less aggressive, more regulatory phenotypes (21). To interrogate this possibility, we performed immune profiling by flow cytometry of pancreatic lymph nodes (pLNs) from mice treated with oral VLX-1005 or vehicle. pLNs are key sites in the initial priming of autoreactive T cells in *NOD* mice (22). Treatment with oral VLX-1005 led to an increase in CD4+Foxp3+ regulatory T cells (Tregs) in the pLNs (**Supplemental Figure 2C**). This effect on Tregs was specific for the pLNs since no changes in Tregs were observed in the spleen after VLX-1005 treatment (**Supplemental Figure 2D**).

### Orally-administered VLX-1005 reduces autoimmune diabetes incidence in female and male NOD.hALOX12 mice

Because 4 weeks of oral VLX-1005 dosing led to improvements in insulitis and reductions in infiltrating T and B cells, we next asked if these alterations lead to prevention or delay of subsequent diabetes development in *NOD.hALOX12* mice. Both female and male mice were administered VLX-1005 via daily oral gavage (30 mg/kg) or vehicle for 4 weeks during the pre-diabetic phase (6-10 weeks of age). Mice were followed for diabetes development (blood glucose ≥250 mg/dl on 2 consecutive days) until 25 weeks of age (**Figure 3A**). At 25 weeks of age, 60% of female mice and 75% of male mice receiving VLX-1005 were protected from diabetes development compared to 25% of female and 50% of male mice receiving vehicle (**Figure 3B-C**). Whereas the preceding studies demonstrate that 12-LOX inhibition with oral VLX-1005 delays the development of diabetes, they do not address if administration of the drug might reverse established diabetes or mitigate hyperglycemia. We allowed female *NOD.hALOX12* mice to develop diabetes (defined as 2 consecutive random-fed blood glucose measurements ≥250 mg/dL), then administered VLX-1005 or vehicle for up to 6 weeks via daily oral gavage or until the mice exhibited signs of physical deterioration from hyperglycemia (loss in body weight, dishevelment) (**Figure 3D**). Notably, we did not observe a reversal in diabetes but did see relative reductions in blood glucose levels in mice treated with VLX-1005 compared to vehicle (**Figure 3E-F**).

**Figure 3:**
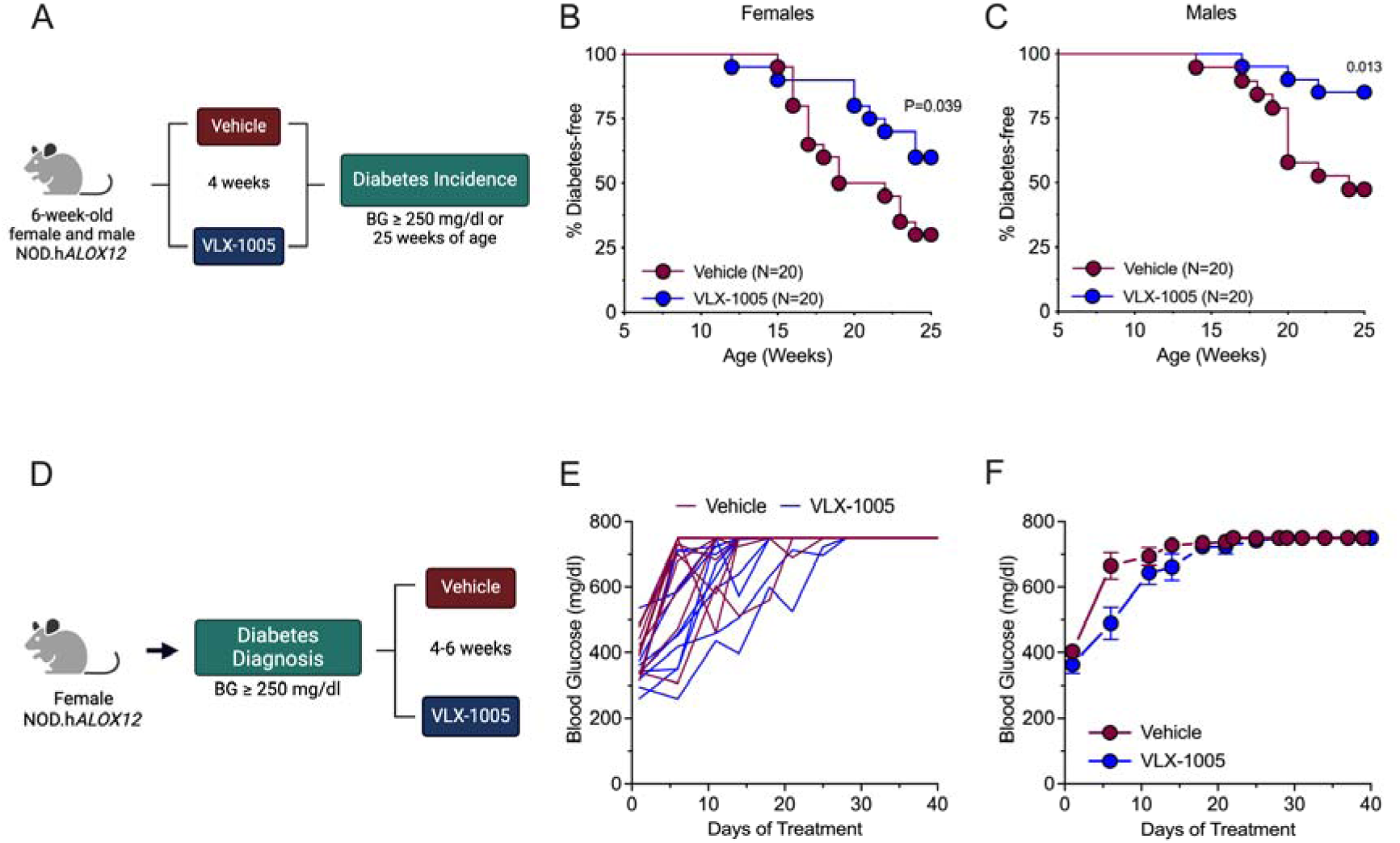
VLX-1005 treatment delays autoimmune diabetes onset in female and male *NOD*.*hALOX12* mice. *NOD.hALOX12* mice (N=20 per group) were treated during the pre-diabetic stage from 6-10 weeks of age or at the time of diabetes development (N=11-12 per group). (**A**) Schematic representation of diabetes prevention experimental design. (**B**) Diabetes incidence in female *NOD*.*hALOX12* mice. (**C**) Diabetes incidence in male *NOD*.*hALOX12* mice. (**D**) Schematic representation of diabetes reversal experimental design. (**E**) Random-fed blood glucose levels in each female mouse. (**F**) Average random-fed blood glucose levels of female mice. Data are presented as mean ±SEM and statistical significance was determined by a Mantel-Cox Log Rank test.

### Orally-administered VLX-1005 reduces islet death and oxidative stress in NOD.hALOX12 mice

12-LOX is primarily present in islets and macrophages, and deletion of the mouse gene (*Alox15*) in either tissue separately was previously shown to reduce diabetes incidence. We, therefore, first asked how treatment with VLX-1005 affects islet cell phenotypes. We first subjected isolated islets from female *NOD.hALOX12* mice treated with vehicle or VLX-1005 to RNA sequencing to identify how islet gene expression might be altered. Principal component analysis of transcriptomics revealed that islets from vehicle- and VLX-1005-treated *NOD.hALOX12* mice clustered separately, suggesting an effect of VLX-1005 treatment on gene expression (**Figure 4A**). Pairwise comparison of gene expression using a false discovery rate (FDR)<0.05 and fold-change (FC)>2 yielded only 189 differentially expressed genes. Instead, a P<0.05 cutoff and FC>2 revealed alteration of 709 genes between vehicle- and VLX-1005-treated *NOD.hALOX12* mice (**Figure 4B**, volcano plot and **Supplemental Table 3** Excel file for full sequencing results). Gene Ontology pathway analysis showed significantly altered pathways related to DNA replication (e.g. *Anp32b, Skp1a, Itfg2, Dmrt1i*), inflammation (NFκB activity) (e.g. *Elf1, Trim75, RNase1, Lmo1, Bcl3, Ptgis, Commd1, Lrrc14, Foxp3*), and G-protein coupled receptor signaling (e.g. *Gpr89, Glp2r*), among others (**Figure 4C**). These pathways suggest responses that may be related to changes in cellular survival in response to VLX-1005. We, therefore, immunostained pancreatic sections for markers of cell death and proliferation in the islet. VLX-1005-treated *NOD.hALOX12* mice exhibited decreased islet cell death as measured by reduced terminal deoxynucleotidyl transferase dUTP nick end labeling (TUNEL) and H2A histone family member X (H2A.X) staining compared to vehicle-treated mice (**Figure 4D**). Additionally, VLX-1005-treated mice demonstrated decreased β cell proliferation, as measured by proliferating cell nuclear antigen (PCNA) immunostaining (**Figure 4D**); reduced PCNA immunostaining was also consistent with the reduction of Ki67 observed in spatial proteomics of the insulin+ area (**Figure 2F** and **Supplemental Figure 2B**). We interpret the reduction in β cell proliferation as a consequence of reduced β cell apoptosis. The alteration in NFκB signaling led us to investigate if markers of oxidative stress were affected since inflammation, oxidative stress, and β cell survival are closely linked (23, 24). We performed immunostaining for the oxidative stress marker, 4-hydroxynonenal (4-HNE) and observed reduced immunostaining in mice treated with VLX-1005 compared to placebo (**Figure 4E**). Consistent with this observation, β cells from VLX-1005-treated animals also displayed an increase in levels of the antioxidant enzyme GPx1 (**Figure 4E**). Collectively, these data are consistent with prior observed effects of reduced inflammation, oxidative stress, and β cell death in β cell-specific deletion of mouse *Alox15* in NOD mice (11).

**Figure 4:**
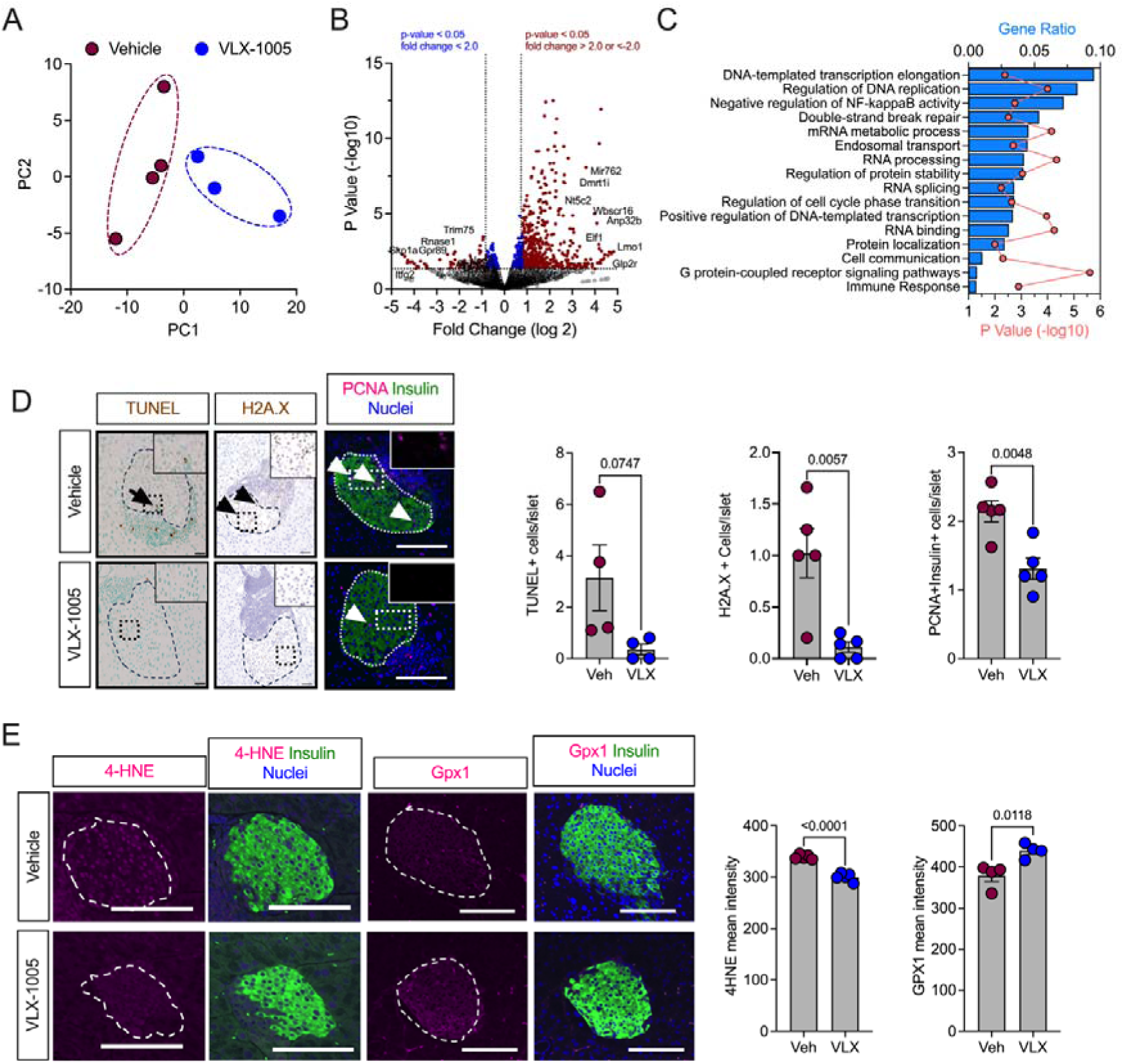
VLX-1005 decreased. β **cell death, proliferation, and oxidative stress in female *NOD*.*hALOX12* mice.** Pancreata or islets were harvested from 10-week-old prediabetic female *NOD*.*hALOX12* mice after 4 weeks of treatment with vehicle or VLX-1005 (N=3-4 per group). (**A**) Principal component analysis plot of RNA-sequencing results from isolated islets of vehicle- or VLX-1005-treated mice. (**B**) Volcano plot of differentially expressed genes. (**C**) Gene ontology pathway analysis of differentially expressed genes. (**D**) Pancreata from mice stained and quantified for TUNEL (brown, *left panels*; black *arrow* indicates positive TUNEL staining within the islet), H2A.X (brown, *middle panels*; black *arrowheads* indicate positive H2A.X staining within the islet), or PCNA (magenta, *right panels*; white *arrowheads* indicate positive PCNA staining within the islet), insulin (green) and nuclei (blue). Each dot represents an individual mouse (N=4-5). Scale bars = 50 μm. (**E**) Pancreata from mice stained and quantified for 4-HNE (magenta, *left panels*), or GPx1 (magenta, *right panels*), and insulin (green) and nuclei (blue). Each dot represents an individual mouse (N=4).Scale bars = 50 μm. Data are presented as mean ±SEM and statistical significance was determined by a two-tailed T-test.

### VLX-1005 alters the proinflammatory macrophage phenotype

Whereas the preceding findings are consistent with improved β cell survival following oral VLX-1005 administration, these studies do not rule out the possibility that the drug directly modifies the phenotype of infiltrating macrophages, which could secondarily affect β cells. Because bulk islet transcriptomics analysis does not resolve gene expression events associated with specific cell types, we isolated bone marrow-derived macrophages (BMDMs) from female *NOD.hALOX12* mice and then performed RNA sequencing in the presence or absence of VLX-1005. BMDMs were unpolarized (“M0”) or polarized to an “M1-like” state (with lipopolysaccharide and IFN-γ) to mimic the inflammatory state that might be observed during T1D pathogenesis. During polarization, BMDMs were treated with vehicle or VLX-1005 (**Figure 5A**). Principal component analysis of transcriptomics revealed that M0 macrophages treated with VLX-1005 co-clustered with vehicle-treated M0 macrophages, suggesting a minimal transcriptional effect of the drug on unpolarized cells (**Figure 5B**). Consistent with this interpretation, only 1% of genes (159 out of 15,888) were significantly altered with VLX-1005 treatment (when using criteria FC>2 and P<0.05) (**Supplemental Table 4** Excel file for full sequencing results). Upon polarization to the M1-like state, a clear rightward shift in the principal component analysis plot was observed with both vehicle and VLX-1005-treated BMDMs and a notable separation was seen between vehicle and VLX-1005 treatment (**Figure 5B**); this finding suggests that the impact of 12-LOX inhibition is more prominent upon a shift to a proinflammatory state of macrophages.

**Figure 5:**
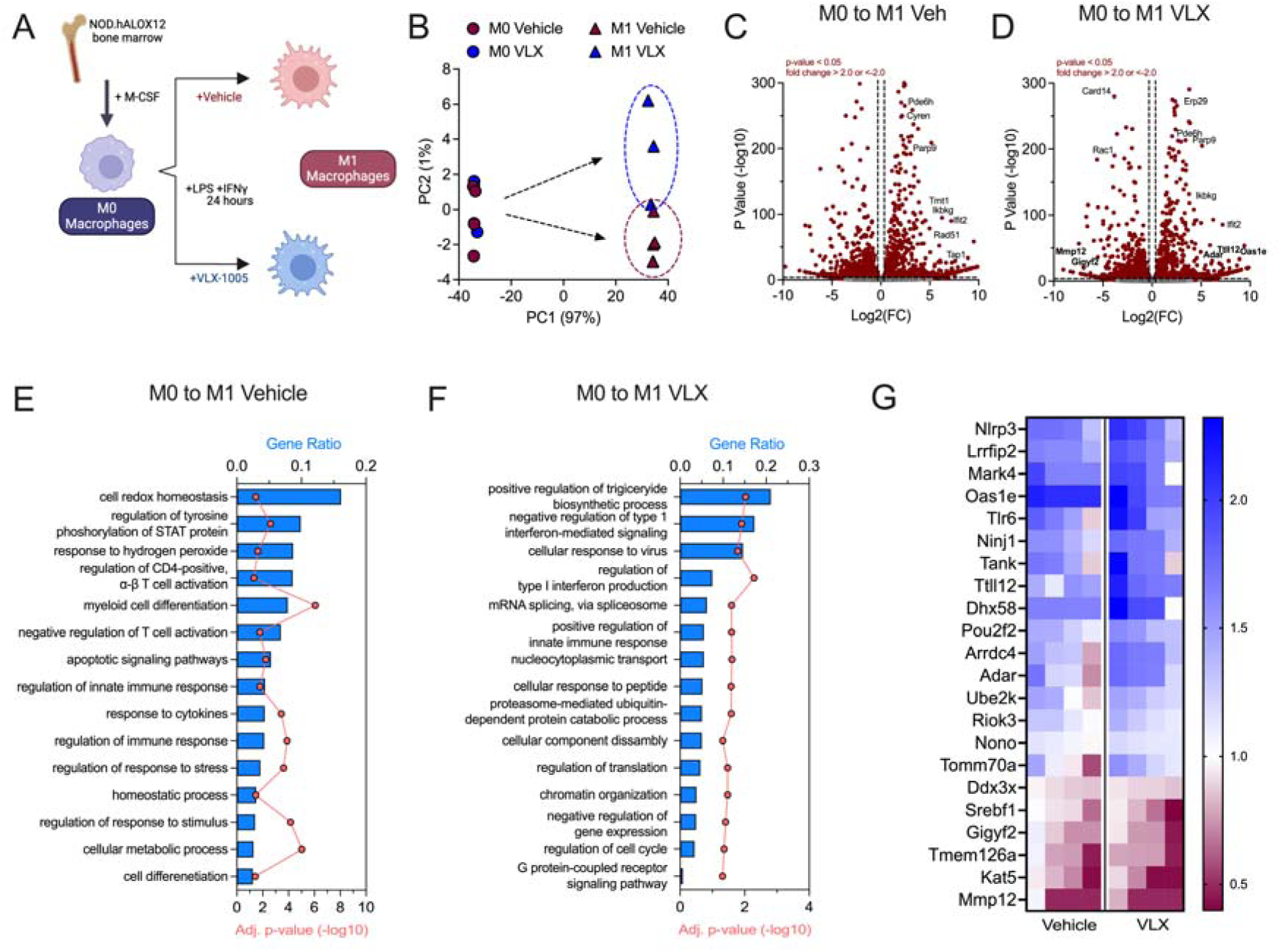
RNA-sequencing analysis of M1-like bone marrow derived macrophages reveals a reduction in the inflammatory response upon VLX-1005 treatment. Bone marrow-derived macrophages (BMDMs) were isolated and polarized to the M1-like state and treated with vehicle or VLX-1005 (10 µM) during polarization. RNA was isolated and sequenced (N=4 per group). (**A**) Schematic of experimental design. (**B**) Principal component analysis plot. (**C**) Volcano plot of differentially expressed genes in M0 and M1-like vehicle-treated macrophages. (**D**) Volcano plot of differentially expressed genes in M0 and M1-like VLX-1005-treated macrophages. (**E**) Gene ontology pathway analysis of differentially expressed genes in M0 vs M1-like vehicle-treated macrophages. (**F**) Gene ontology pathway analysis of differentially expressed genes in M0 vs M1-like VLX-1005-treated macrophages. (**G**) Heatmap of significantly altered interferon-related genes. Columns represent sequencing results from each sample (N=4 per group). Numbers on the heatmap scale indicate fold change compared to M0 macrophages.

We next interrogated the gene expression events associated with the M0 to M1 transition in both vehicle and VLX-1005-treated BMDMs. Most genes altered (FC>2 and P<0.05) in this transition (2467) were common between vehicle and VLX-1005 treatment (**Figure 5C-D**, **Supplemental Figure 3A,** and **Supplemental Table 4** Excel file). These common genes mapped to pathways related to cytokine-mediated signaling, T cell activation, and antigen processing and presentation (**Supplemental Figure 3B**). An additional 507 genes were significantly altered in vehicle-treated cells, and 459 additional genes were significantly altered with VLX-1005 treatment (**Supplemental Figure 3A**). The 507 genes altered with vehicle treatment mapped to GO pathways related to the M1 polarization phenotype (myeloid cell differentiation, immune response, response to cytokines) and pathways related to oxidative stress (cell redox homeostasis, response to hydrogen peroxides) (**Figure 5E**). These pathways were not identified in the genes that were differentially expressed during VLX-1005 treatment. GO pathway analysis of the 459 genes altered with VLX-1005 showed particularly significant alterations in pathways related to modification of the interferon response (**Figure 5F**). Next, we looked specifically at the genes that mapped to interferon pathways (**Figure 5G**), which revealed that VLX-1005 augmented significantly (P<0.05 by T-test) the magnitude of the gene changes compared to vehicle treatment. Notable genes, whose directional changes are known to counter the interferon response included *Oas1e* (25), *Ttll12* (26), and *Adar* (27) (all upregulated compared to control-treated M1 macrophages), and *Gigyf2* (28) and *Mmp12* (29) (correspondingly downregulated) (**Figure 5G**). These data suggest effects of VLX-1005 that may lead to reduced macrophage interferon signaling.

## DISCUSSION

To date, the adaptive immune system has remained the primary focus for the development of therapeutics aimed at preventing or reversing T1D. Notwithstanding the utility of agents such as anti-CD3 monoclonal antibodies in preserving β cell function (30) or delaying T1D development (7), there has been impetus in the research community to develop therapeutics that target other cell types that contribute to T1D development (31), including innate immune cells and β cells. A multi-targeted approach is expected to aid in better disease modification and result in more durable and broadly applicable therapy (32). In this respect, 12/15-LOX (in mice) is a particularly appealing target since it is active in both macrophages and β cells and contributes to the development of inflammatory disorders, including insulin resistance, atherosclerosis, and T1D (for review, see (9)). The deletion of *Alox15*, specifically in either myeloid cells or islet β cells, proved sufficient to delay/prevent T1D in NOD mice (2, 11). Evidence for 12-LOX contributions to T1D pathogenesis identifies this enzyme as an attractive target in human disease. 12-LOX is elevated in β cells of autoantibody-positive, pre-T1D individuals and in residual β cells of individuals with established T1D (12). A pro-inflammatory product derived from the 12-LOX (and mouse 12/15-LOX)-mediated metabolism of arachidonic acid is 12-HETE, an eicosanoid that either directly or indirectly (through G-protein-coupled receptors (33–36)) augments reactive oxygen species generation and endoplasmic reticulum stress in macrophages and islets (37, 38). Notably, levels of 12-HETE were shown to be elevated in the circulation of youth and adults with new-onset T1D (compared to healthy controls and those with established T1D) (39).

Considering the biological contributions of the 12-LOX enzymes (mouse 12/15-LOX and human 12-LOX) to T1D and other inflammatory disorders, the development of enzyme inhibitors offers an attractive approach to disease modification. Inhibition of 12/15-LOX using ML351 demonstrated promising outcomes in NOD mice, with reductions in insulitis and improvements in glucose homeostasis (13). An inhibitor that showed mouse and human cross-species reactivity, ML127, unfortunately also displayed evidence of off-target cytotoxicity (13). By contrast, VLX-1005 (a.k.a. ML355) is a potent inhibitor (IC_50_ ∼300 nM) of human 12-LOX (14) that has shown efficacy in reversing the insulin secretory defects of cytokine-treated human islets in vitro without evidence of cytotoxicity (17); however, the benefit of this inhibitor in disease states in vivo has remained speculative. To address the translational challenge of testing 12-LOX inhibitors in preclinical disease models in vivo, we developed a human gene replacement mouse model on both the C57BL/6J and NOD autoimmune diabetes backgrounds. The utility of this model as a platform for 12-LOX inhibitor testing was confirmed in our multiple low-dose STZ studies, which showed that VLX-1005 administration to *B6.ALOX12* mice precluded hyperglycemia, whereas wild-type controls developed hyperglycemia over time. STZ is a toxin whose full effects involve communication between β cells and macrophages (19), and our findings with systemic administration of VLX-1005 are consistent with similar STZ studies in mice harboring the global deletion of *Alox15* (18)—collectively suggesting that human *ALOX12* gene replacement mice respond appropriately to a 12-LOX inhibitor and that the human *ALOX12* gene can sufficiently replace the functionality of the mouse *Alox15* gene.

In recent studies, our group showed that loss of the mouse *Alox15* gene in either myeloid cells or islet β cells could protect animals from the development of autoimmune diabetes on the NOD background (2, 11). The pathological phenotypes of these animals were similar, with reductions in islet invasion by T cells, B cells, and myeloid cells and a characteristic increase in PD-L1 in either macrophages or β cells. PD-L1 is an immune checkpoint protein whose interaction with its receptor PD-1 on adaptive immune cells leads to a more immune-suppressive response (21). In our studies using *NOD.ALOX12* mice, we found similar responses to VLX-1005 treatment, with striking reductions in both innate and adaptive immune cell infiltration into the islets and a notable increase in PD-L1 in the insulitic (immune cell) component. We interpret these latter findings to suggest that the effect of systemically administered VLX-1005 may be greater in macrophages than in β cells. The reductions in markers of β cell proliferation, death, and oxidative stress that we observed with VLX-1005 treatment may also be reflective of this preferential effect on macrophages since these responses are otherwise characteristic of the effects of cytokines on β cells (40). Our RNA-Seq studies of isolated NOD mouse BMDMs polarized to the proinflammatory M1 state support this contention, as VLX-1005 treatment resulted in reductions in the interferon response. Our studies open the possibility that VLX-1005 may be useful in autoimmune diabetes resulting from PD-L1 or PD-1 blockade (checkpoint inhibitor therapy) often used to treat cancers (41).

Some limitations to our study should be acknowledged. First, because our mouse model replaces the mouse *Alox15* with the human *ALOX12* globally, we cannot be certain that the effects we observed are exclusively related to the inhibition of the human enzyme in only macrophages and β cells; a recent study suggested that mouse *Alox15* may contribute to pro-resolving functions of Tregs (42). It remains unclear, therefore, if the loss of *Alox15* globally in our mouse model might have affected Treg function or the function of other cell types that have low levels of *Alox15* expression. In this respect, our mouse model may have limited utility in other disease states where *ALOX12* might not fully replace the function of *Alox15.* Moreover, we cannot rule out the possibility that *Alox15* and *ALOX12* mediate distinct pathways that nevertheless still allow for shared phenotypes. For example, our RNA sequencing studies of macrophages treated with VLX-1005 reveal distinct reductions in interferon-mediated responses but such effects have not been directly explored as a consequence of *Alox15* in mice. Second, our studies do not fully address the timing and duration of VLX-1005 treatment. We only treated mice for a 4-week period (6-10 weeks of age); it is possible that a longer duration of treatment might have yielded even more robust T1D prevention outcomes. Therefore, the timing and duration of human treatment may require further investigation in our preclinical model. Limitations regarding timing and duration may also apply to diabetes “reversal” studies, in which we observed a reduction in glycemia but no reversal in disease. We cannot rule out the possibility that the effect of VLX-1005 to suppress β cell proliferation may have prevented a more robust outcome in this case. Finally, our study does not assess the potential for 12-LOX inhibition in combination with other immunomodulatory agents. Whereas the relatively modest impact of VLX-1005 treatment in the diabetes reversal studies in *NOD.ALOX12* mice might suggest diminishing returns on the treatment of humans at the time of disease diagnosis, this effect may be amplified in the presence of T or B cell blockade.

Despite these limitations, our studies identify a new platform on which to study a class of LOX inhibitors for their utility in ameliorating human autoimmune diabetes. Our human gene replacement mouse model demonstrates a functional equivalence between mouse *Alox15* and human *ALOX12* in the context of T1D since the whole-body replacement of the mouse gene with the human (under the mouse upstream control elements) preserves islet pathology and the frequency of diabetes incidence in NOD mice. Therefore, beyond its utility to test inhibitors of human 12-LOX, our mouse model also provides a platform to interrogate the cause-effect relationship of human 12-LOX in T1D and possibly other inflammatory diseases in vivo.

## MATERIALS and METHODS

### Sex as a biological variable

For *C57BL/6J* mice, our study examined only male mice, because comparative data for these mice in the literature are primarily from males. For *NOD* mice, our study mostly examined females, because type 1 diabetes in the *NOD* strain is more frequently observed in females. However, some data from *NOD* male mice are included that parallel those seen in females, suggesting that the effects observed in females may be relevant to males.

### Animals

Male and female *C57BL/6J* mice and *NOD/ShiLTJ* mice were procured from the Jackson Laboratory. All mice were kept under pathogen free housing conditions with standard light:dark (12:12 h) cycles and fed ad lib normal chow. To generate a humanized *ALOX12* mouse model, the coding region of the mouse *Alox15* gene was replaced with the coding region of the human *ALOX12* gene while retaining all the mouse regulatory elements (**Figure 1A**) (mice were generated by a contract to Ingenious Targeting Laboratory). Targeted iTL BF1 (*C57BL/6* FLP) embryonic stem cells were microinjected into *Balb/c* blastocysts. Chimeras with high percentage of black coat color resulting from this procedure were then mated to *C57BL/6J* wildtype mice to generate germline neo-deleted mice. The following primers were used to genotype the mice: 5’-TCTGATCTGTGTATGCCTGTGTGTGG-3’ (forward) and 5’-TTCCAAGGAAAAAGGCATGGTTTCTGAGG-3’ (reverse). These primers generate a 478 bp band for wildtype and 581 bp band for knock-in mice (**Supplemental Figure 1A**).

Human *ALOX12* alleles were introgressed onto both the *C57BL/6J* and *NOD.ShiLT/J* mouse backgrounds using a speed congenics approach based on microsatellite genotyping at The Jackson Laboratory. Genome scanning was also performed at The Jackson Laboratory to confirm successful backcrossing onto the *C57BL/6J* and *NOD.ShiLT/J* mouse background (*B6*.*hALOX12* and *NOD*.*hALOX12*; **Supplemental Table 1** excel file). Body mass was measured by EchoMRI.

Intraperitoneal glucose tolerance tests (IPGTT) were performed in mice after overnight fasting (16 h). Mice were interperitoneally injected with glucose at a dose of 1 or 2 g/kg body weight and blood glucose levels were measured at specific time points: 0, 10, 20, 30, 60, 90, and 120 minutes after glucose injection using an AlphaTrak glucometer.

### Formulation of VLX-1005

VLX-1005 was obtained from Veralox Therapeutics Inc (Frederick, MD). A VLX-1005 spray-dried dispersion (for oral administration) was prepared by dissolving VLX-1005 and HPMC-E3 in a 90:10 w/w mixture of tetrahydrofuran and water to attain a total solids concentration of 5% w/w. 1575 g of solution was then spray dried using a Buchi B-290 laboratory spray dryer. The yield after spray drying was 67.8 g. The collected material was further dried in an oven at 40 °C under vacuum to remove residual tetrahydrofuran.

### Pharmacokinetic Analysis and Lipidomics

Following intraperitoneal injection or oral gavage, VLX-1005 was quantified in plasma using high-performance liquid chromatography-tandem mass spectrometry (Triple Quad 6500+; Sciex) after separation by HPLC (Column: Agilient Poroshell 120 EC-C18; HPLC: Shimadzu DGU-405). Pharmacokinetic parameters for VLX-1005 were estimated by non-compartmental model using WinNonlin 8.3. The bioavailability (F%) was calculated as the following: AUC_last_-PO/AUCINF-PO > 80%: F=(AUCINF-PO*DoseIV)/(mean AUCINF-IV*DosePO). Lipidomics on serum samples was performed by the New York Medical College Lipidomics Core using a Shimadzu LC-MS/MS 8050 system equipped with a UHPLC and auto-sampler.

### Streptozotocin (STZ) Induction

Male *C57BL/6J* and *B6*.*hALOX12* mice (8-10 weeks of age) were injected with either vehicle (0.5% methylcellulose) or 30 mg/kg/day of VLX-1005 by intraperitoneal injection for 15 days: 5 days prior to the start of multiple low-dose STZ (55 mg/kg/day; 5 consecutive days), 5 days during STZ treatment, and 5 days post STZ injections. Male *B6.hALOX12* mice (8-10 weeks of age) were injected with either vehicle (0.5% methylcellulose) or 30 mg/kg/day of VLX-1005 by oral gavage (PO) for 11 days: 3 days prior to the start of multiple low dose STZ (55 mg/kg/day; 5 consecutive days), 5 days during STZ treatment, and 3 days post STZ injections. Random-fed glucose levels were measured by tail snip using a glucometer (AlphaTrak), and mice were followed for 20 days post-STZ injections. IPGTT was performed on day 4 post-STZ treatments after overnight fasting. At the end of each study, mice were euthanized, and pancreas and blood samples were collected.

### Diabetes Incidence and Treatment

Both male and female *NOD*.*hALOX12* mice were given either vehicle or 30 mg/kg/day VLX-1005 (PO) for 4 weeks in the pre-diabetic stage (6-10 weeks of age) and then followed for diabetes incidence until 25 weeks of age or until diabetes diagnosis. Diabetes incidence was determined by observing two consecutive blood glucose values greater than 250 mg/dL. At the end of each study, mice were euthanized, and pancreas and blood samples were collected.

For diabetes treatment studies, female *NOD*.*hALOX12* mice were followed for random-fed blood glucose from 12-20 weeks of age. At diabetes incidence (two consecutive blood glucose values greater than 250 mg/dL), mice were administered 30 mg/kg/day VLX-1005 SDD or vehicle for up to 6 weeks via daily oral gavage or until the mice exhibited signs of physical deterioration from hyperglycemia (loss in body weight, dishevelment). At the end of each study, mice were euthanized, and pancreas and blood samples were collected.

### Islet and Macrophage Isolation

Islets were isolated from *NOD*.*hALOX12* mice with either vehicle or 30 mg/kg/day of VLX-1005treatment using collagenase digestion. Briefly, collagenase was injected into the pancreatic bile duct to digest the connective tissue and release pancreatic cells (43). A Histopaque-HBSS gradient was applied to the dissociated pancreas and centrifuged at 900 *xg* for 18 min. The isolated islets were cultured in RPMI medium. The collected islets were handpicked and allowed to recover overnight before processing. RNA was isolated for use in RNA sequencing or quantitative PCR.

Bone marrow-derived macrophages (BMDMs) were isolated from 8-week-old *NOD*.*hALOX12* mice as described previously (2). The isolated BMDMs were cultured for 7 days in complete medium (RPMI containing 10% FBS, 10 mM HEPES, and 100 U/ml penicillin/ streptomycin) supplemented with 10 ng/ml M-CSF. On day 7 of culture, the BMDMs were pretreated with either vehicle (0.1% DMSO) or 10 μM VLX-1005. After 1 h pretreatment, the BMDMs were further stimulated with 10 ng/ml LPS and 25 ng/mL IFN-γ for 18 h for M1-like polarization. RNA was isolated and used for sequencing.

### RNA Isolation and Quantitative PCR

RNA was isolated from mouse tissues and macrophages using an RNeasy Mini® Kit from Qiagen. The isolated RNA was used to synthesize cDNA using a High-Capacity cDNA Reverse Transcription kit (Applied Biosystems) according to manufacturer’s instructions. Quantitative PCR was performed using a Bio-Rad CFX Opus with a predesigned Taqman® assay probe for human and mouse genes: human *ALOX12*: Hs00167524_m1; mouse *Alox15*: Mm00507789_m1; mouse *Actb*: Mm01205647_m1 (Invitrogen). The relative gene expression levels were calculated using the comparative threshold cycle value (Ct) and normalized to *Actb*.

### Immunostaining, ***β*** Cell Mass, and Insulitis Scoring

Pancreatic tissues were fixed using 4% paraformaldehyde. After fixation, the tissues were embedded in paraffin and sectioned with a thickness of 5 μm. Three sections per mouse were used for analysis, with each section being spaced 100 μm apart. Tissue sections were immunostained with anti-insulin (ProteinTech; 15848-1-AP; 1:200), anti-glucagon (Abcam; ab92517; 1:200), anti-12-LOX (Thermo Fisher; PA5-26020; 1:200), anti-12/15-LOX (Abcam; ab80221; 1:200), anti-CD3 (Abcam; ab16669; 1:200), anti-F4/80 (Sigma: D2S9R; 1:150), anti-MAC2 (Thermo Fisher; EbioM3/38; 1:200) and anti-H2A.X (Cell Signaling Technology; 9718s; 1:200) primary antibodies followed by conjugated anti-rabbit Ig (Vector Laboratories) secondary antibody. A DAB (3,3’-diaminobenzedine) Peroxidase Substrate Kit from Vector Laboratories was used for detection. After immunostaining, the tissue sections were counterstained with hematoxylin (Sigma). Images were acquired using a BZ-X810 fluorescence microscope (Keyence) and β/α cell mass was quantified by insulin+ or glucagon+ area and whole pancreas area. Insulitis score reflects the degree of immune cell infiltration within pancreatic islets. The score system used as follow: 1 = no insulitis, 2 = infiltrate <50% circumference, 3 = infiltrate >50% circumference, 4 = infiltration within islet. Data are shown as the average insulitis score per mouse.

For immunofluorescence staining, pancreatic sections were stained with the following antibodies: anti-insulin (Dako IR002; 1:4), anti-glucagon (Santa Cruz; sc514592; 1:50), anti-B220 (Biolegend; 03201; 1:100), anti-CD3 (Abcam; ab16669; 1:200), anti-PCNA (Santa Cruz; sc-7907; 1:100), anti-4HNE (Abcam; ab46545; 1:200), and anti-GPx1(Santa Cruz; sc-22145; 1:100). Highly cross-adsorbed Alexa Fluor secondary antibodies (ThermoFisher) were used at a dilution of 1:500. Tissue sections were stained with DAPI (ThermoFisher) to label cell nuclei. The Nikon A1 confocal microscopy was used to capture images. CellProfiler v4.1 software was used for image analysis.

### TUNEL Staining

Terminal deoxynucleotidyl transferase dUTP nick end labelling (TUNEL) assay was used to determine β cell death in pancreatic islets. The assay was performed according to the protocol provided by the manufacturer (Abcam) and HRP-DAB chemistry was used for detection. Two sections, spaced 100 μm apart, were used for each mouse. Images were captured using a BZ-X810 fluorescence microscope system (Keyence). The number of TUNEL+ cells was assessed manually per islet.

### NanoString Spatial Proteomics

Paraffin embedded pancreata were used for NanoString spatial proteomics analysis. Tissues were stained with morphology markers: AF-647 conjugated insulin (Cell Signaling; 9008s; 1:400) and nuclei marker (SYTO13). Tissues were hybridized using a pre-validated mouse GeoMx Immune cell panel (NanoString; GMX-PROCONCT-MICP) comprising of the following markers: PD-1, CD11c, CD8a, PanCk, MHC II, CD19, CTLA4, SMA, CD11b, CD3e, Fibronectin, Ki-67, CD4, GZMB, F4/80, CD45, PD-L1; housekeeping genes: Histone H3, S6, GAPDH; and IgG antibodies: Rb IgG, Rat IgG2a, and Rat IgG2b for background subtraction. All markers were conjugated to unique UV-photocleavable oligos for indexing. At least 5-6 islets with insulitis were chosen as regions of interest (ROI) per mouse based on the morphology markers (insulin and nuclei). The ROIs were segmented into insulitic region and insulin+ region for each islet. Oligos from the segmented ROIs were photocleaved, collected in a 96-well plate, and reads were counted using nCounter (NanoString). Scaling was performed to normalize for any differences in tissue surface area and depth. After scaling, reads were normalized to housekeeping markers and background was subtracted using IgG markers.

### Flow Cytometry

Spleen and pancreatic lymph nodes were harvested, homogenized, and passed through a 70 μm strainer to obtain a single cell suspension. Cell pellets were resuspended in red blood cell (RBC) lysing solution to remove red blood cells. 2.5*10^5-1^*10^6^ cells per condition were incubated with blocking solution (eBioscience; 14-0161-86) containing anti-mouse CD16/CD32 to block the Fc receptors for 20 min on ice. The following surface markers were used — CD4-FITC (BioLegend; 100510; 1:100), CD8-PerCP-Cy5.5 (Biolegend; 100734; 1:100), CD19-AF700 (Biolegend; 152414; 1:100). Following incubation of surface antibodies, cells were washed with stain buffer and then permeabilized using fix/perm buffer (BD #554722) before intracellular staining. The following intracellular antibodies were used FoxP3-AF647 (BD; 560401; 1:100), IFNγ-PE (BD; 554412; 1:50), and IL17a-APCCY7 (BD; 560821; 1:50). Cells were analyzed on the Attune NxT Flow Cytometer (Thermo Fisher). Data were analyzed by FlowJo software (BD Biosciences).

### RNA Sequencing

RNA extraction was performed using RLT Buffer, according to the manufacturer’s instructions (Qiagen). Samples were submitted for library generation and sequencing by the University of Chicago sequencing core using a NovaSeq 6000^®^ (Illumina). Data was analyzed using Galaxy (https://usegalaxy.org/). Reads were aligned to the Mus musculus genome build mm10 using HISAT2. Individual sample reads were quantified using HTseq-count and normalized using DESeq2. DEseq2 was also used to calculate fold changes and P-values and to perform optional covariate correction. Gene ontology (GO) was used for pathway analysis.

### Statistical Methods

All data are represented as mean *±* SEM. When comparing more than two conditions, one-way ANOVA was performed. Tukey’s post-hoc test or Dunnett’s post-hoc test was used to determine specific differences between individual group means. When comparing only two conditions, two-tailed student’s t-test was performed. Mantel-Cox log-rank test was specifically used for analyzing the NOD diabetes incidence experiments. Data analyses were performed using the GraphPad Prism 10 software. The differences were considered statistically significant at a p value <0.05.

### Study Approval

All experiments involving mice were performed at the University of Chicago and the procedures were conducted according to protocols approved by the University of Chicago Institutional Animal Care and Use Committee (Chicago, IL).

## Data Availability

The islet RNA sequencing data have been uploaded to the Gene Expression Omnibus (https://www.ncbi.nlm.nih.gov/geo/) with accession number GSE272668. The BMDM sequencing data have been uploaded to the Gene Expression Omnibus with accession number GSE272687. Values for all data points in graphs are reported in the Supporting Data Values file.

## ACKNOWLEDGEMENTS

This work was supported in part by National Institutes of Health grants R03 TR003381 (to SAT and RGM), R41 DK122917 (to RGM and DJM), R01 DK10558 (to RGM), U01 DK127786 (to RGM), T32 AI153020 (to JRE), an investigator-initiated award from Veralox Therapeutics (to SAT and RGM), a Chicago Biomedical Consortium Director’s Award (to RGM), and Breakthrough T1D postdoctoral fellowship (3-PDF-2023-1326-A-N) and Diabetes Research Connection awards (both to CM). This study utilized Diabetes Center core resources supported by National Institutes of Health grant P30 DK020595 (to the University of Chicago) and utilized services of the University of Chicago Histology and Genomics Cores.

## AUTHOR CONTRIBUTIONS

JLN, DJM, MBB, SAT, and RGM conceptualized the research; TN, CM, JRE, JEW, AC, KF, SP, JBN, and SAT performed investigation; SAT and RGM provided project supervision; TN, SCM, SAT, and RGM wrote the original draft; all authors contributed to discussion, edited the manuscript, and approved the final version of the manuscript.

## DECLARATION OF INTERESTS

RGM and SAT received an investigator-initiated award from Veralox Therapeutics. RGM serves on the Scientific Advisory Board for Veralox Therapeutics. DJM and MBB are Veralox Therapeutics employees.

## Notes

### Summary of Updates

Title change, small changes to figures (ie removing of bold lettering).

