## Supplemental Data for "12-Lipoxygenase inhibition delays onset of autoimmune diabetes in human gene replacement mice"

**Supplemental Figure 1: Comparison of *B6*.*hALOX12* to *C57BL/6J* mice and *NOD*.*hALOX12* to *NOD.ShiltJ* mice at 10 weeks of age.** (**A**) Genotyping results of *B6.hALOX12* mice. (**B**) Body weight. (**C**) Fat mass. (**D**) Lean mass. (**E**) Random-fed blood glucose. (**F**) IPGTT and AUC (*right panel*) of 8 week old male *B6.hALOX12* compared to *C57BL/6J* mice. (**G**) Pancreata from the mice indicated stained for glucagon (magenta), insulin (green) and nuclei (blue) (left panels); insulin (brown, middle panels); or glucagon (brown, right panels). Scale bars = 50 μm. (**H**) β cell mass, each dot represents an individual mouse. (**I**) α cell mass, each dot represents an individual mouse. (**J**) Pancreata from female *NOD* and *NOD.hALOX12* mice stained for 12/15-LOX (magenta), insulin (green), and nuclei (blue) (left panels). Scale bars = 20 μm. (**K**) Pancreata from female *NOD* and *NOD.hALOX12* mice stained for 12-LOX (magenta), insulin (green), and nuclei (blue) (left panels). Scale bars = 20 μm. (**L**) Pancreata from female *NOD* and *NOD.hALOX12* mice stained for CD3 (magenta), B220 (teal), insulin (white), and nuclei (blue) (left panels) or insulin (brown, right panels). Scale bars = 50 μm. (**M**) Insulitis score, each dot represents data from an individual mouse. Data are presented as mean ±SEM and statistical significance was determined by a two-tailed T-test.

**Supplemental Figure 2: Spatial proteomics analysis and flow cytometry analysis of *NOD.hALOX12* mice treated with vehicle or VLX-1005.** Female *NOD.hALOX12* mice were treated for 4 weeks with either vehicle or VLX-1005. Nanostring® spatial proteomics were performed in the pancreas in regions of interest that included the peri-islet insulitic area (**A**) or the intra-islet area (**B**) for the indicated markers. Each dot represents a different region of interest. Pancreatic lymph nodes (**C**) or spleen (**D**) were isolated from treated mice and subjected to flow cytometry for the indicated markers. Each dot represents data from a single mouse. Data are presented as mean ±SEM and statistical analysis was performed using a two-tailed T-test.

**Supplemental Figure 3: RNA-sequencing analysis of M1-like bone marrow derived macrophages (BMDM).** BMDMs were isolated and polarized to the M1-like state and treated with vehicle or VLX-1005 (10 µM) during polarization. RNA was isolated and sequenced. (A) Differential gene expression identified between vehicle- and VLX-1005-treated macrophages during the M0 to M1 transition phase. (B) Gene ontology pathway analysis of the common differentially expressed genes.

**Supplemental Table 2: Lipidomics results of non-12-lipoxygenase products from serum of mice treated with vehicle or VLX-1005 for 4 weeks.** Data are presented as mean ±SEM and statistical analysis was performed using a two-tailed T-test.

|  | **Vehicle** | **30 mg/kg VLX-1005 SDD** | **p-value** |
| --- | --- | --- | --- |
| **5-HETE** | 96.1± 35.3 | 16.0 ± 2.5 | 0.053 |
| **11(12)-EET** | 18.1 ± 8.1 | 2.2 ± 0.4 | 0.083 |
| **14,15-DHET** | 12.6 ± 4.4 | 5.4 ± 0.7 | 0.141 |
| **11,12-DHET** | 6.4 ± 2.3 | 4.0 ± 0.5 | 0.350 |
| **14(15)-EET** | 16.1 ± 7.8 | 3.6 ± 0.6 | 0.110 |
| **8(9)-EET** | 24.8 ± 11.1 | 3.8 ± 1.0 | 0.096 |
| **5(6)-EET** | 140.7 ± 55.3 | 24.2 ± 5.3 | 0.070 |
| **18-HETE** | 3.0 ± 1.5 | 1.7 ± 0.4 | 0.433 |
| **TX-B2** | 12.4 ± 2.7 | 5.5 ± 2.2 | 0.086 |
| **6-keto-PGF1a** | 39.2 ± 11.8 | 1.1 ± 0.6 | 0.012 |
| **Leukotriene-B4** | 1.5 ± 0.7 | 0.1 ± 0.1 | 0.080 |
| **5-oxo-ETE** | 12.2 ± 3.7 | 3.9 ± 0.8 | 0.058 |
| **13-HDHA** | 19.1 ± 7.0 | 3.4 ± 0.9 | 0.056 |
